# Cell-type deconvolution of bulk RNA-Seq from kidney using opensource bioinformatic tools

**DOI:** 10.1101/2023.02.13.528258

**Authors:** Angelica M. Riojas, Kimberly D. Spradling-Reeves, Clinton L. Christensen, Shannan Hall-Ursone, Laura A. Cox

## Abstract

Traditional bulk RNA-Seq pipelines do not assess cell-type composition within heterogeneous tissues. Therefore, it is difficult to determine whether conflicting findings among samples or datasets are the result of biological differences or technical differences due to variation in sample collections. This report provides a user-friendly, open source method to assess cell-type composition in bulk RNA-Seq datasets for heterogeneous tissues using published single cell (sc)RNA-Seq data as a reference. As an example, we apply the method to analysis of kidney cortex bulk RNA-Seq data from female (N=8) and male (N=9) baboons to assess whether observed transcriptome sex differences are biological or technical, i.e., variation due to ultrasound guided biopsy collections. We found cell-type composition was not statistically different in female versus male transcriptomes based on expression of 274 kidney cell-type specific transcripts, indicating differences in gene expression are not due to sampling differences. This method of cell-type composition analysis is recommended for providing rigor in analysis of bulk RNA-Seq datasets from complex tissues. It is clear that with reduced costs, more analyses will be done using scRNA-Seq; however, the approach described here is relevant for data mining and meta analyses of the thousands of bulk RNA-Seq data archived in the NCBI GEO public database.

**Author Summary:** This method, which provides a simple method for assessing sampling biases in bulk RNA-Seq datasets with evaluation of cell-type composition, will aid researchers in assessing whether bulk RNA-Seq from different studies of the same heterogeneous tissue are comparable. The additional layer of information can help determine if differential gene expression observed is biological or technical, i.e., cell composition variation among study samples. The described method uses publicly available bioinformatics resources and does not require coding expertise or high-capacity computational processing. Development of tools accessible to scientists without computing expertise will contribute to greater rigor and reproducibility for bioinformatic analyses of transcriptome data.

## Introduction

Bulk RNA-Seq is a sensitive and robust method for identifying transcriptional changes in tissues, but it’s often difficult to assess what cell types are represented in transcriptome data. Standard computational pipelines for bulk RNA-Seq data analysis do not typically assess cell-type composition within heterogeneous tissues(1–4). This gap leads to uncertainty when seemingly similar tissue samples are compared and show significantly different gene expression profiles across studies, bringing reproducibility of findings into question, i.e., are observed transcriptional differences due to biological differences or sampling differences among samples(5–8). In order to accurately compare RNA-Seq data among samples collected from heterogeneous tissues, cell-type composition must be addressed. Therefore, there is a need to develop user friendly, open source and simple method to identify cell types across samples and studies.

While some methods allow for cell type selection for bulk RNA-Seq during sample preparation such as tissue microdissection and flow cytometry, the additional processing requires collection of greater amounts of tissue than simple bulk RNA-Seq(9–13). Single cell (sc) or single nuclei (sn) RNA-Seq can be a more challenging albeit rewarding task, and may be performed using flow cytometry, cell selection by antibodies, or microfluidic devices to separate cell types (9–14). These methods can be limited by expense, amount of sample available, sample preparation, and sample heterogeneity. This is especially challenging for more complex tissues such as the kidney with significant cell-type heterogeneity, including shape and size, and where only small tissue biopsies are available(5,14,15).

Quality control (QC) assessment of bulk RNA-Seq data does not typically include assessment of cell-type composition, rather QC is limited to RNA quality(16,17). This consists of assessing RNA integrity prior to library preparation. Library insert fragment length are assessed after library preparation(1–3,18). Quality of sequencing chemistries are measured by Phred scores which calculate the error probability by evaluating peak shape and resolution at each nucleotide - with current sequencing instruments this is obtained during each round of amplification in the sequencer (1–3,18). Overall dataset quality after sequencing is then gauged by average read lengths and percent alignment to a reference genome (1–3,18). While this is informative, it does not provide information on the cell types present in the samples.

Single cell references use a list of previously identified gene markers based on cell-type specific gene and protein expression to identify cell types(8,14,19,20). These analyses have not only validated previously identified cell types in tissues, but also revealed new cell types and importantly new cell-type specific markers(8,14,19,20). We can now use these genes to perform cell-type composition analysis on bulk RNA-Seq datasets without the need to run a parallel scRNA-Seq experiment from a subset of samples included in the experiment. While methods are available for cell deconvolution, most require some coding skill, subscriptions to specialized hosting packages, and sequence data derived from paired bulk RNA-Seq and scRNA-Seq experiments(18,21,22). These methods are often inaccessible to scientists with limited bioinformatics training.

We describe an easy-to-use protocol for those with limited computational background to process a previously published single cell dataset for creation of a reference cell-type expression list, and then annotate a bulk RNA-Seq dataset using Microsoft Office, which is commonly available at minimal expense in universities and other workplaces. This method can easily be adapted to R or other software with a simple script.

We also provide an example using this method for cell type composition analysis of kidney cortex biopsy samples obtained from female and male baboons. Baboons (*Papio hamadryas*) are genetically and physiologically similar to humans and have been used to study the role of the kidney in many complex diseases(21–30). The structures and substructures within the kidney are quite heterogeneous and to date, more than 40 different cell types have been identified among mammals(14,20). We performed bulk RNA-Seq on ultrasound guided kidney cortex biopsies (females, n=8; males, n=9). We generated a reference list of renal cell-type specific genes using data from Clark *et al*. which included 144 genes from 38 cell types(20). We determined that cell-type representation was similar between females and males, indicating that observed differences in gene expression were not due to sampling bias. This example demonstrates the utility of the method for assessing contribution of multiple cell types to bulk RNA-Seq expression for evaluation of biological versus technical variation.

## Methods

### Ethics, study design, and data collection

We utilized a cohort of baboons (females: n=8, age 17.88 ± 1.35 years; males: n=9, age 8.10 ± 2.38 years)(29–31) (*Papio hamadryas*; Taxonomy ID 9557) maintained as part of the pedigreed baboon colony at SNPRC, located on the campus of Texas Biomedical Research Institute, San Antonio, Texas. All animal procedures were reviewed and approved by Texas Biomedical Research Institute’s Institutional Animal Care and Use Committee protocols (IACUC). SNPRC facilities at the Texas Biomedical Research Institute and animal use programs are accredited by Association for Assessment and Accreditation of Laboratory Animal Care International (AAALAC), operate according to all National Institutes of Health (NIH) and U.S. Department of Agriculture (USDA) guidelines, and are directed by veterinarians (DVM). All animal care decisions were made by the SNPRC veterinarians. Daily enrichment was provided by the SNPRC veterinary staff and behavioral staff in accordance with AAALAC, NIH, and USDA guidelines. The KINSHIP program from Pedigree Database System (PEDSYS v. 2.0) was used with the Stevens-Boyce algorithm to calculate kinship coefficients to ensure similar degrees of relatedness among all animals(29–31).

### Kidney biopsy collection

All animals were preoperatively treated with ketorolac (5 mg/kg), sedated with ketamine (10mg/kg, IM), and maintained on isoflurane (1.3-3.0%) anesthesia throughout kidney biopsy. Kidney biopsies were collected via ultrasound guidance from the cortex region of the left kidney and immediately placed into cooled 1.5 mL tubes then flash frozen in liquid nitrogen. Animals were monitored during collections and throughout recovery. Postoperative care included observation for swelling at the surgical sight and monitoring until animal was conscious. Buprenorphine (0.2 mg/kg, SQ) was administered for post-operative pain relief immediately once animals were awake. Buprenorphine (0.2 mg/kg, SQ) was administered as needed 24-48 hours after surgery(29–31).

### Diet and housing

The baboons were raised and maintained on a standard monkey chow diet (high complex carbohydrates; low fat (“Monkey Diet 15%/5LEO,” LabDiet, PMI Nutrition International) prior to study initiation. Animals were acclimated to cages for at least 8 weeks prior to study. Animals had free access to 500 g chow *ad libitum* in individual cages during feeding times. Female baboons were housed in an outdoor social group with one vasectomized male (not on study) to provide full social and physical activity. Females were run into individual cages during feeding times. Males were housed individually prior to study start for 6 weeks(29,30,32).

### RNA isolation, library preparation, and sequencing

RNA was isolated from kidney biopsy samples using the Direct-zol RNA Miniprep Plus Kit (Zymo Research; R2051). Tissue was homogenized in 600 μL of TRI reagent (Zymo Research; R2051) using a BeadBeater (Biospec Products) for 3 × 30 sec with RNA purification according to the Direct-zol RNA Miniprep Plus Kit instructions. RNA was quantified by Qubit RNA BR Assay Kit (Invitrogen; Q10210), and RNA integrity was determined with an RNA ScreenTape Kit on the TapeStation 2200 system (Agilent; 5067-5576, 5067-5578, 5067-5577) and only RIN scores of 8 or higher were used for sequencing. cDNA libraries were prepared using the KAPA Stranded RNA-Seq Kit (Roche; 07962142001). Libraries were quantified using a KAPA Library Quantification Kit (Roche; KR0405), and quality was determined with a DNA ScreenTape Kit on the TapeStation 2200 system (Agilent; 5067-5582, 5067-5583). Sequencing was done using a HiSeq 2500 (Illumina).

### RNA-Seq data processing

Female and male transcript data were processed together in Partek^®^ Flow^®^ (Partek^®^, Inc.) with a minimum read length of 25 and minimum Phred score of 330 30. Adapter sequences were trimmed with the resulting average read length above 90 bases in each group. Reads were aligned to the baboon genome (papAnu2.0; March 2012). Reads were normalized by TMM+1 and filtered for genes expressed an average of less than 1 across all samples. The list of baboon kidney bulk RNA-Seq quality transcripts for all samples was saved as a .csv file. The list included official gene symbols and abundance values for each gene in each sample.

### Assessing bulk RNA-Seq sample cell-type composition

The list of transcripts that passed quality filters was annotated to identify kidney-specific cell types by the following steps. A reference list of kidney cell-type specific transcripts was derived from previously published rat whole kidney scRNA-Seq data which included genes specifically expressed in one of 43 kidney cell types(20). For annotation of our bulk RNA-Seq data, each rat kidney cell type gene list was downloaded and combined into a single rat kidney cell type list in MS Excel. The list included columns with official gene symbol, whole kidney expression value, and kidney cell type annotation. In excel, the tool “Conditional Formatting” was used to identify and highlight duplicate values, sorted according to color, and highlighted (duplicate) values were deleted. The resulting list included genes that were highly expressed and uniquely expressed in only one kidney cell type. The rat kidney cell type list was saved in a .csv format.

The rat kidney cell type list and the baboon kidney bulk RNA-Seq quality transcript list were uploaded onto the Galaxy server (usegalaxy.org)(18). The Galaxy tool join/subtract/group> join two datasets was used to merge the two lists according to the respective columns with official gene symbols. The merged output .txt file was downloaded and converted to a .csv file. The resulting merged file provides cell-type annotation for genes in the baboon bulk RNA-Seq file that were included in the rat scRNA-Seq kidney cell type list (n=274).

Data, the 274 cell-type specific transcripts were assessed. Multiple unpaired t-tests with FDR two-stage setup (Benjamini, Krieger, and Yekutieli) to adjust for multiple testing were performed in GraphPad Prism version 9.5.0 for Mac, (GraphPad Software). Unbiased gene expression analysis by PCA was performed on normalized transcripts with ellipsoids representative of 2 standard deviations away from the mean using Partek Genomics Suite (Partek^®^, Inc.). Data are expressed as mean, mean ± SD, or coefficient of variation and were statistically significant if the FDR adjusted p-value < 0.05.

## Results

We identified a group of baboons with low degree of relatedness representing diverse genetic backgrounds from the SNPRC colony (Additional File 1 tab S1). We performed a standard workflow for quality filtering of bulk RNA-Seq data prior to comparison of females and males to identify differentially expressed genes(30,32). We assessed pre-alignment total reads, average read length, and average Phred quality score (Fig 1A, 1B, 1C). Post alignment quality control measures included average depth, average coverage depth, and percent reads aligned to the reference transcriptome (Fig 1D, 1E, 1F). Average read length significantly differed by Welch’s t-tests as expected from differences in library preparation (p value=0.0117). The percentage alignment was also significantly different (p value=0.0434), which may be explained by sex differences.

**Fig 1.**
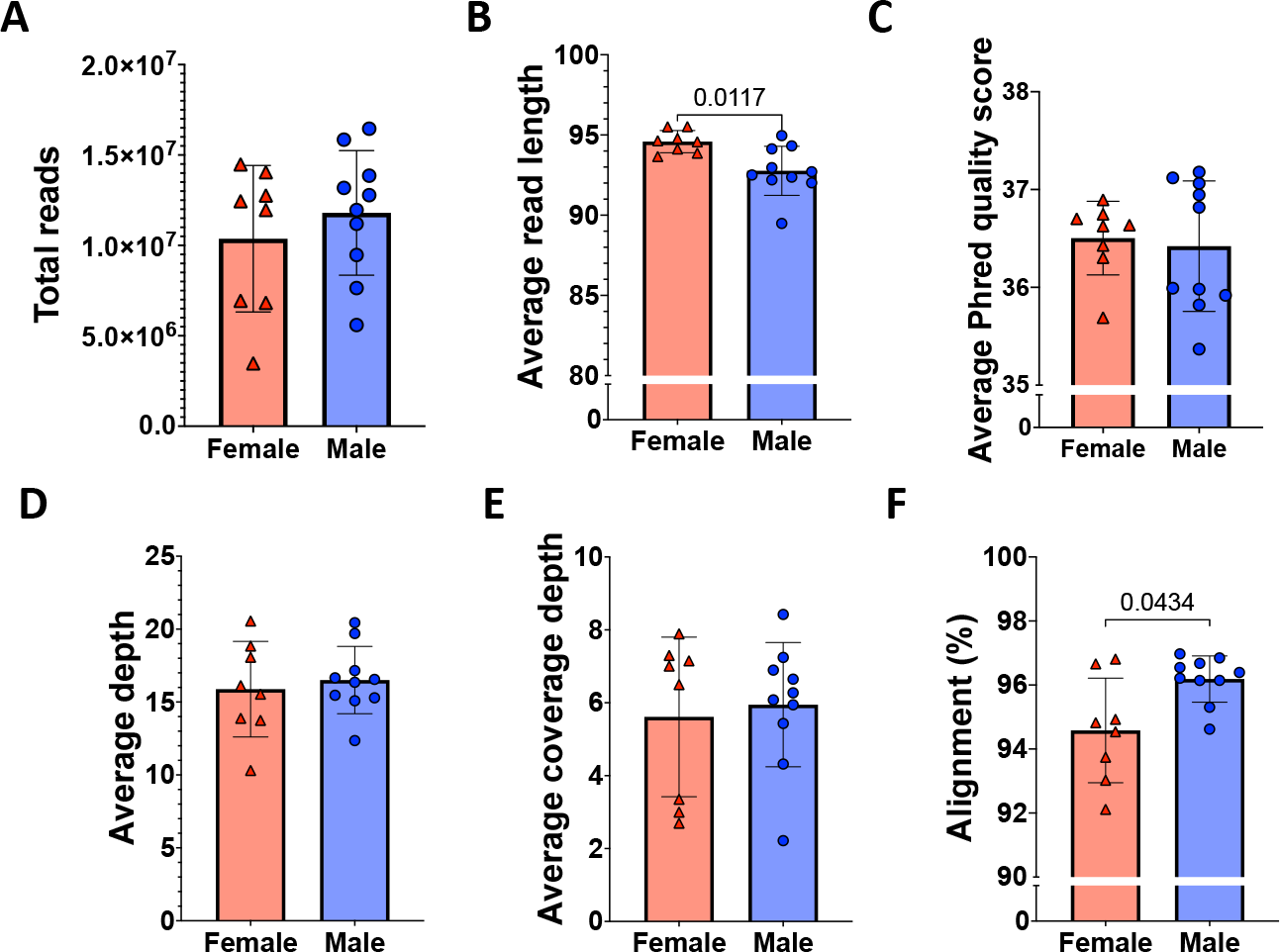
Sequence data quality and alignment of female and male kidney cortex biopsy reads. Reads were assessed for quality prior to downstream analysis in female (n=8; red triangles) and male (n=9; blue circles) based on: A. Total reads, B. Average read length, C. Average Phred quality score, D. Average depth, E. Average coverage depth, and F. Alignment (%). Unpaired t-test with Welch’s correction was performed comparing males to females in each panel; p-values listed above bars if significance was <0.05. Individual values are shown, as well as the mean represented by columns, with error bars indicating standard deviation.

### Cell type-specific gene expression in kidney cortex biopsies

We assessed cell-type composition of the baboon transcriptome (Additional File 1 tab S2). Fig 2 shows the 40 highest expressed transcripts. Kidney cortex transcripts included 38 of the 43 kidney cell types previously characterized in rat by Clark *et al*.(20) among all female and male baboon samples. The baboon transcript list contained 274 transcripts encoded by 144 cell-type specific genes, i.e., some genes included more than one transcript isoform (Fig 3, Additional File 1 tab S3). No significant differences were observed across 274 transcripts by multiple unpaired t-tests. Only one gene, *SLC18A2* found in neural axon cells (a low abundance cell type), was expressed significantly higher in males compared to females (p value=0.000024).

**Fig 2.**
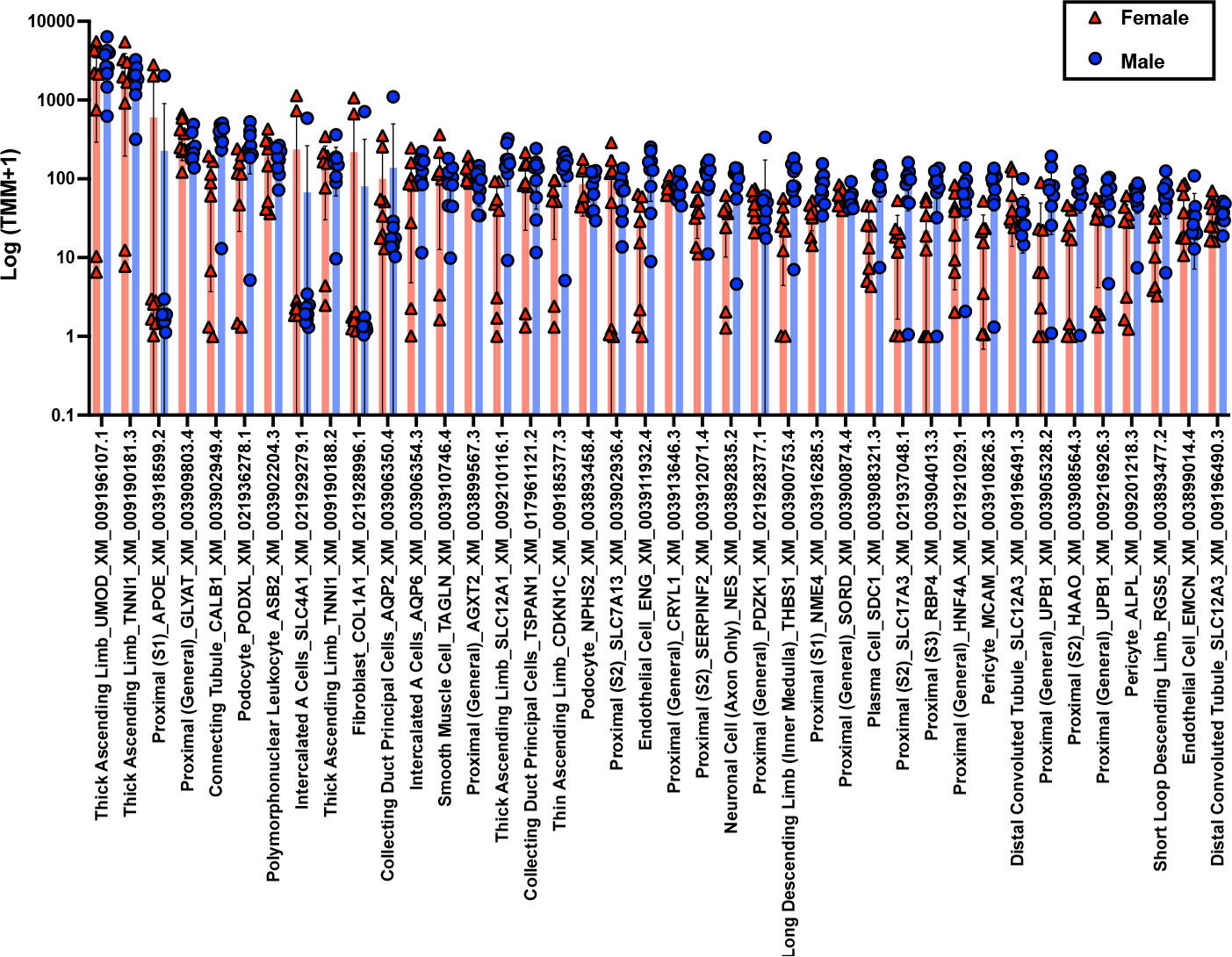
Top 40 highest expressed cell-type specific transcripts in female and male kidney cortex biopsies. The y-axis indicates abundance, and the x-axis indicates cell type, official gene symbol, and NCBI transcript ID for each transcript. Individual values for females (n=8) are denoted by red triangles and males (n=9) by blue circles. The mean values of transcripts are represented by columns with error bars indicating standard deviations. Multiple unpaired t-tests with FDR threshold of 1.00% and two-stage setup (Benjamini, Krieger, and Yekutieli) were used. No significant differences were observed between males and females.

**Fig 3.**
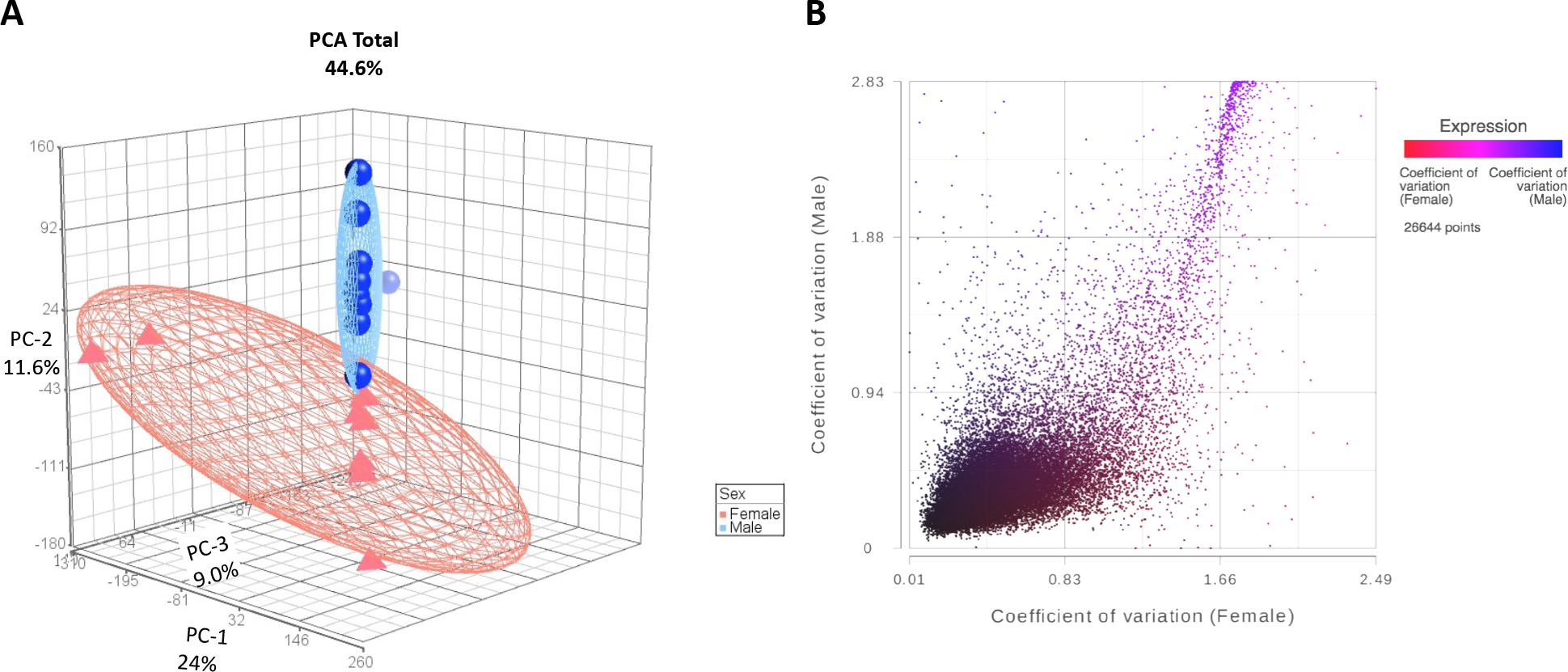
Variation in gene expression of female (n=8) and male (n=9) kidney cortex biopsies. A. Principal component analysis of female and male kidney cortex transcriptomes. Top principal components totaled 44.6% and represented each axis as follows: 1=24%, 2=11.6%, and 3=9.03%. Each female transcriptome is represented as a red triangle and each male transcriptome as a blue sphere. Ellipsoids denote 2 standard deviations from the mean for each group. B. Coefficient of variation in 26,644 female and male kidney cortex biopsy transcripts. Each point demonstrates a single transcript with the coefficient of variation value in females on the x axis (n=8) and males (n=9) on the y axis. The color of each point indicates if the transcript expression level (TMM+1) is greater in females (red) or males (blue).

### Sex differences in baboon kidney cortex transcriptome

PCA of all quality transcripts in female and male baboon samples revealed the top principal components of 24%, 11.6%, and 9.03%, which account for 44.6% variation in the samples, with two clusters revealing sex-specific profiles (Fig 3A). Coefficient of variation for transcripts in females and males showed sex differences in gene expression variation among individuals and in the distribution of transcriptome variation (Fig 3B, Additional file 1 tab S4). A subset of genes, *FAM19A3, LEP, NOS1*, were only found to be expressed in female baboon kidney cortex samples.

### Applications

Many bulk RNA-Seq studies have been performed to date, and rarely is it clear whether gene expression differences are due to tissue sampling (technical) or biological differences. Profiling cell composition strengthens results by ensuring greater rigor and reproducibility. By using gene expression that is exclusively expressed in one cell type, it is reasonable to infer that the cell type is present. Most of the currently available methods to deconvolute cell types in bulk RNA-Seq data require hosting packages and may be inaccessible to scientists with limited bioinformatics training. We present a simple method for cell type deconvolution.

This new method ultimately saves time, samples, and money by leveraging existing datasets as a reference. Moreover, collaborative groups often have many people involved from sample collection to final analysis, and cell type profiling provides an additional layer of QC for the person doing downstream data analysis. This technique can also be applied to older datasets, proteomic and metabolomic datasets, and aid in determining study similarity related to tissue and cell-type sampling(33,34). It is worth noting that published studies that appear to show conflicting results may benefit from cell-type deconvolution to assess technical versus biological differences. In addition, this approach is relevant to meta-analyses of publicly available bulk RNA-Seq data.

## Discussion

We performed bulk RNA-Seq on kidney cortex biopsies to assess sex differences in the transcriptome and used a simple cell type deconvolution method to determine whether observed differences were technical or biological. Our results showed consistency in kidney biopsy cell composition among samples indicating the sex differences were biological. This method of assessing sampling consistency of bulk RNA-Seq datasets utilizing an existing scRNA-Seq dataset as a reference with a “computationally light” annotation pipeline, is especially useful for bulk RNA-Seq studies with heterogeneous cell types. Inclusion of these steps provides an additional level of rigor for transcript data quality assessment. Use of official gene symbols, in which orthologous genes use the same symbol, allows cross species comparison to leverage the extensive publicly available scRNA-Seq datasets primarily generated from rodent studies(18,27,35–37). The method is relevant to assessing technical variation across studies with conflicting findings, and for potential meta-analyses of publicly available bulk RNA-Seq data.

Limitations: While this method does not tell us the number of cells present for each cell type, it consistently reveals the presence of individual cell types allowing comparison among samples(18,30,35,37).

## Data Sharing Statement—Large biological datasets

RNA-Seq data are available at NCBI GEO GSE166295.

## Funding and support

This investigation was conducted in facilities constructed with support from Research Facilities Improvement Program Grant Numbers C06 RR015456 and C06 RR013556 from the National Center for Research Resources (NCRR), National Institutes of Health (NIH). Other resources used in this study were supported by NIH grant 5 R01 HL68180, TL1 TR002647, SNPRC grant P51 OD011133 from the Office of Research Infrastructure Programs (ORIP), NIH.

## Disclosure statement

There are no disclosures.

## Competing Interest Statement

There are no competing interests with any of the authors involved in this work.

## Acknowledgements

None

## Notes

### Competing Interest Statement

The authors have declared no competing interest.

## References

1. Yalamanchili HK, Wan Y, Liu Z. Data Analysis Pipeline for RNA-seq Experiments: From Differential Expression to Cryptic Splicing. Curr Protoc Bioinformatics. 2017 Sep 13;59(1).

2. Sanchis P, Lavignolle R, Abbate M, Lage-Vickers S, Vazquez E, Cotignola J, et al. Analysis workflow of publicly available RNA-sequencing datasets. STAR Protoc. 2021 Jun;2(2):100478.

3. Conesa A, Madrigal P, Tarazona S, Gomez-Cabrero D, Cervera A, McPherson A, et al. A survey of best practices for RNA-seq data analysis. Genome Biol. 2016 Dec 26;17(1):13.

4. Vieth B, Parekh S, Ziegenhain C, Enard W, Hellmann I. A systematic evaluation of single cell RNA-seq analysis pipelines. Nat Commun. 2019 Oct 11;10(1):4667.

5. Conway BR, O’Sullivan ED, Cairns C, O’Sullivan J, Simpson DJ, Salzano A, et al. Kidney Single-Cell Atlas Reveals Myeloid Heterogeneity in Progression and Regression of Kidney Disease. Journal of the American Society of Nephrology. 2020 Dec;31(12):2833–54.

6. Noureen N, Ye Z, Chen Y, Wang X, Zheng S. Signature-scoring methods developed for bulk samples are not adequate for cancer single-cell RNA sequencing data. Elife. 2022 Feb 25;11.

7. Hegenbarth JC, Lezzoche G, de Windt LJ, Stoll M. Perspectives on Bulk-Tissue RNA Sequencing and Single-Cell RNA Sequencing for Cardiac Transcriptomics. Frontiers in Molecular Medicine. 2022 Feb 22;2.

8. Dong M, Thennavan A, Urrutia E, Li Y, Perou CM, Zou F, et al. SCDC: bulk gene expression deconvolution by multiple single-cell RNA sequencing references. Brief Bioinform. 2021 Jan 18;22(1):416–27.

9. Zhang X, Wang Z, Zhang C, Li Y, Lu S, Steffens S, et al. Laser Capture Microdissection–Based mRNA Expression Microarrays and Single-Cell RNA Sequencing in Atherosclerosis Research. In 2022. p. 715–26.

10. Zilionis R, Nainys J, Veres A, Savova V, Zemmour D, Klein AM, et al. Single-cell barcoding and sequencing using droplet microfluidics. Nat Protoc. 2017 Jan 8;12(1):44–3.

11. Zhou W min, Yan Y yan, Guo Q ru, Ji H, Wang H, Xu T tian, et al. Microfluidics applications for high-throughput single cell sequencing. J Nanobiotechnology. 2021 Dec 11;19(1):312.

12. Nguyen QH, Pervolarakis N, Nee K, Kessenbrock K. Experimental Considerations for Single-Cell RNA Sequencing Approaches. Front Cell Dev Biol. 2018 Sep 4;6.

13. Kashima Y, Sakamoto Y, Kaneko K, Seki M, Suzuki Y, Suzuki A. Single-cell sequencing techniques from individual to multiomics analyses. Exp Mol Med. 2020 Sep 15;52(9):1419–27.

14. Park J, Shrestha R, Qiu C, Kondo A, Huang S, Werth M, et al. Single-cell transcriptomics of the mouse kidney reveals potential cellular targets of kidney disease. Science (1979) [Internet]. 2018/04/07. 2018;360(6390):758–63. Available from: https://www.ncbi.nlm.nih.gov/pubmed/29622724

15. Muto Y, Wilson PC, Ledru N, Wu H, Dimke H, Waikar SS, et al. Single cell transcriptional and chromatin accessibility profiling redefine cellular heterogeneity in the adult human kidney. Nat Commun. 2021 Apr 13;12(1):2190.

16. Hong M, Tao S, Zhang L, Diao LT, Huang X, Huang S, et al. RNA sequencing: new technologies and applications in cancer research. J Hematol Oncol. 2020 Dec 4;13(1):166.

17. Kukurba KR, Montgomery SB. RNA Sequencing and Analysis. Cold Spring Harb Protoc. 2015 Nov;2015(11):pdb.top084970.

18. Afgan E, Nekrutenko A, Grüning BA, Blankenberg D, Goecks J, Schatz MC, et al. The Galaxy platform for accessible, reproducible and collaborative biomedical analyses: 2022 update. Nucleic Acids Res. 2022 Jul 5;50(W1):W345–51.

19. Freytag S, Tian L, Lönnstedt I, Ng M, Bahlo M. Comparison of clustering tools in R for medium-sized 10x Genomics single-cell RNA-sequencing data. F1000Res. 2018 Dec 19;7:1297.

20. Clark JZ, Chen L, Chou CL, Jung HJ, Lee JW, Knepper MA. Representation and relative abundance of cell-type selective markers in whole-kidney RNA-Seq data. Kidney Int. 2019;95(4):787–96.

21. Bronikowski AM, Alberts SC, Altmann J, Packer C, Carey KD, Tatar M. The aging baboon: Comparative demography in a non-human primate. Proceedings of the National Academy of Sciences. 2002 Jul 9;99(14).

22. Rogers J, Raveendran M, Harris RA, Mailund T, Leppälä K, Athanasiadis G, et al. The comparative genomics and complex population history of Papio baboons. Sci Adv [Internet]. 2019 Jan 30;5(1):eaau6947. Available from: https://advances.sciencemag.org/lookup/doi/10.1126/sciadv.aau6947

23. Kammerer CM, Cox LA, Mahaney MC, Rogers J, Shade RE. Sodium-Lithium Countertransport Activity Is Linked to Chromosome 5 in Baboons. Hypertension [Internet]. 2001 Feb;37(2):398–402. Available from: https://www.ahajournals.org/doi/10.1161/01.HYP.37.2.398

24. McGill HC, McMahan C a., Kruski a. W, Mott GE. Relationship of lipoprotein cholesterol concentrations to experimental atherosclerosis in baboons. Arterioscler Thromb Vasc Biol [Internet]. 1981;1(1):3–12. Available from: http://atvb.ahajournals.org/cgi/doi/10.1161/01.ATV.1.1.3

25. Cox LA, Comuzzie AG, Havill LM, Karere GM, Spradling KD, Mahaney MC, et al. Baboons as a Model to Study Genetics and Epigenetics of Human Disease. ILAR J [Internet]. 2013 Jan 1;54(2):106–21. Available from: https://academic.oup.com/ilarjournal/article-lookup/doi/10.1093/ilar/ilt038

26. Carey D, Kammerer CM, Shade RE, Rice KS, McGill Jr. HC. Selective breeding to develop lines of baboons with high and low blood pressure. Hypertension [Internet]. 1993/06/01. 1993;21(6 Pt 2):1076–9. Available from: https://www.ncbi.nlm.nih.gov/pubmed/8505095

27. Benjamin EJ, Virani SS, Callaway CW, Chamberlain AM, Chang AR, Cheng S, et al. A custom rat and baboon hypertension gene array to compare experimental models. Science (1979) [Internet]. 2012/01/10. 2012;93(1):99–110. Available from: https://www.ncbi.nlm.nih.gov/pubmed/28793269

28. Spradling KD, Glenn JP, Garcia R, Shade RE, Cox LA. The baboon kidney transcriptome: analysis of transcript sequence, splice variants, and abundance. PLoS One [Internet]. 2013/05/03. 2013;8(4):e57563. Available from: https://www.ncbi.nlm.nih.gov/pubmed/23637735

29. Riojas AM, Spradling-Reeves KD, Shade RE, Puppala SR, Christensen CL, Birnbaum S, et al. Sex Differences in Blood Pressure and the Kidney Cortex Transcriptome in Nonhuman Primates. Research Gate. 2021 Oct 28;(PREPRINT).

30. Riojas AM, Reeves KD, Shade RE, Puppala SR, Christensen CL, Birnbaum S, et al. Blood pressure and the kidney cortex transcriptome response to high-sodium diet challenge in female nonhuman primates. Physiol Genomics. 2022 Nov 1;54(11):443–54.

31. Spradling-Reeves KD, Shade RE, Haywood JR, Cox LA. Primate response to angiotensin infusion and high sodium intake differ by sodium lithium countertransport phenotype. J Am Soc Hypertens [Internet]. 2017/02/28. 2017;11(3):178–84. Available from: https://www.ncbi.nlm.nih.gov/pubmed/28238630

32. Riojas, A M; Spradling-Reeves, K D; Shade, R E; Puppala, S R; Christensen, C L; Birnbaum, S; Glenn, J P; Hall-Ursone, S; Cox LA. Sex Differences in Blood Pressure and the Kidney Cortex Transcriptome in Nonhuman Primates. preprint [Internet]. Available from: https://bio.tools/pedsys

33. Sigdel TK, Piehowski PD, Roy S, Liberto J, Hansen JR, Swensen AC, et al. Near-Single-Cell Proteomics Profiling of the Proximal Tubular and Glomerulus of the Normal Human Kidney. Front Med (Lausanne). 2020 Sep 17;7.

34. Wang G, Heijs B, Kostidis S, Rietjens RGJ, Koning M, Yuan L, et al. Spatial dynamic metabolomics identifies metabolic cell fate trajectories in human kidney differentiation. Cell Stem Cell. 2022 Nov;29(11):1580–1593.e7.

35. Fan J, Lyu Y, Zhang Q, Wang X, Li M, Xiao R. MuSiC2: cell-type deconvolution for multi-condition bulk RNA-seq data. Brief Bioinform. 2022 Nov 19;23(6).

36. Wu H, Uchimura K, Donnelly EL, Kirita Y, Morris SA, Humphreys BD. Comparative Analysis and Refinement of Human PSC-Derived Kidney Organoid Differentiation with Single-Cell Transcriptomics. Cell Stem Cell [Internet]. 2018/11/20. 2018;23(6):869–881e8. Available from: https://www.ncbi.nlm.nih.gov/pubmed/30449713

37. Avila Cobos F, Alquicira-Hernandez J, Powell JE, Mestdagh P, de Preter K. Benchmarking of cell type deconvolution pipelines for transcriptomics data. Nat Commun. 2020 Nov 6;11(1):5650.

